# CosTaL: An Accurate and Scalable Graph-Based Clustering Algorithm for High-Dimensional Single-Cell Data Analysis

**DOI:** 10.1101/2022.11.10.516044

**Authors:** Yijia Li, Jonathan Nguyen, David Anastasiu, Edgar A. Arriaga

**Affiliations:** Department of Biochemistry, Molecular Biology, and Biophysics, University of Minnesota, Minneapolis, Minnesota; Department of Computer Science and Engineering, Santa Clara University, Santa Clara, California; Department of Chemistry, University of Minnesota, Minneapolis, Minnesota

**Keywords:** Clustering, Mass Cytometry, Flow Cytometry, Single-cell RNA sequencing, Graph-based clustering, *k* nearest neighbors

## Abstract

With the aim of analyzing large-sized multidimensional single-cell datasets, we are describing our method for Cosine-based Tanimoto similarity-refined graph for community detection using Leiden’s algorithm (CosTaL). As a graph-based clustering method, CosTaL transforms the cells with high-dimensional features into a weighted k-nearest-neighbor (kNN) graph. The cells are represented by the vertices of the graph, while an edge between two vertices in the graph represents the close relatedness between the two similar cells. Specifically, CosTaL builds an exact kNN graph using cosine similarity and uses the Tanimoto coefficient as the refining strategy to re-weight the edges in order to improve the effectiveness of clustering. We demonstrate that CosTaL generally achieves equivalent or higher effectiveness scores on seven benchmark cytometry datasets and six single-cell RNA-sequencing datasets using six different evaluation metrics, compared with other state-of-the-art graph-based clustering methods, including PhenoGraph, Scanpy, and PARC. CosTaL is also the most efficient algorithm on large datasets, suggesting that CosTaL generally has better scalability than the other methods, which is beneficial for large-scale analysis.

## Introduction

Cells can be classified into different types according to their intrinsic heterogeneity in proteins, nucleotides, and other metabolites. The classification of cells facilitates the understanding of relationships between cell identities and their functions in disease, aging, and other biological models. In the past several years, multiparametric single-cell profiling methods, such as Mass Cytometry (MC) and single-cell RNA sequencing (scRNA-seq), have significantly improved the characterization of single cells, allowing researchers to depict different types of cells precisely from a complex population (1, 2).

Nowadays, cytometry methods such as MC allow measuring up to more than 40 parameters per cell, while the scRNA-seq method generally yields more than 20,000 features during a single measurement (3, 4). The manual gating method can be used to identify cell types based on the signature of the features using histograms and bivariate plots. However, gating graphically is relatively subjective and time-consuming.

For sequencing data, due to the large number of parameters contained in the data, exhaustive manual gating approaches are not feasible. Thus, researchers often resort to automated unsupervised clustering methods in order to characterize cell types (5, 6). A growing trend of generating larger scale single-cell data is expected as a result of the rapid development and widespread application of single-cell techniques in recent years. This makes it increasingly important for clustering algorithms to allow users to process large-sized multiparametric datasets accurately and promptly.

In order to deal with large datasets with both high dimensionality and a large number of cells, a straightforward strategy is down-sampling, which selects only a portion of the total cells for analysis. This has been done by algorithms like SPADE (7). However, down-sampling has the potential to overlook rare populations. On the other hand, if no down-sampling is conducted, scalability becomes an issue for analyzing large datasets. Based on this perspective, clustering algorithms with high complexity like SNN-Clip, and SC3 are unsuitable for large datasets as they require increased computational workloads and execution time (8). Additionally, clustering algorithms should be extensible so that emerging analysis pipelines or analysis protocols for new single-cell techniques can use them directly. Many algorithms developed specifically for single-cell platforms have limitations in extending the strategy to other studies employing different methods. X-shift, for example, is designed for cytometry datasets and cannot be applied to scRNA-seq data (7, 9). BackSPIN is another algorithm that is only applicable to scRNA-seq datasets rather than cytometry datasets. Among all the existing algorithms that are both scalable and extensible, FlowSOM, PhenoGraph, Seurat, and Scanpy are the most commonly used methods (10). FlowSOM, although being extremely fast, requires a parameter *k* as the desired cluster number. In most cases, *k* should be determined by the user, as studies have demonstrated that automatic estimation within the algorithm is not always helpful (5). Also, FlowSOM has an over-partitioning problem that generates too many small clusters, making it hard to interpret the clustering results, especially without prior knowledge of the populations (5, 11). On the other hand, PhenoGraph, Seurat, and Scanpy are algorithms that share the strategy of transforming the high-dimensional cell data into a graph to represent the similarity relationships between the cells. Similarly, PARC is another efficient graph-based clustering algorithm that specializes in detecting rare populations (12).

With the aim of developing a more efficient, scalable, and extensible clustering algorithm for large single-cell datasets, we formulate CosTaL, a graph-based single-cell clustering framework. In comparison with other single-cell clustering methods, CosTaL has superior efficiency and competitive effectiveness when tested on a total of 13 benchmark datasets. When clustering scRNA-seq datasets using CosTaL, one should note that no normalization, scaling, or Principal Components Analysis (PCA) transformations are required, which significantly reduces the effort involved in parameter selection as well as increases the overall clustering efficienc

## Methods

### A. Description of the CosTaL algorithm

Current methods for profiling single cells capture many features, such as those represented by different fluorescence channels in flow cytometry (FC). Each one of those features with unique signal intensities, corresponds to one dimension in a highdimensional Euclidean space. Similar types of cells should share a similar pattern of distribution for most of the features, resulting in a dense region in the high-dimensional space which can be used to identify the cell clusters.

Across all the existing unsupervised clustering algorithms, Jaccard-Louvain methods, as exemplified in PhenoGraph, PARC, Seurat, and Scanpy, are the most widely used strategies (6, 10). These methods are effective due to the basis that building a k-Nearest Neighbor (kNN) graph is an efficient means of extracting high-dimensional distributions of the cells and preserving their relatedness within dense areas (6, 10, 13–18). In a kNN graph, cells are converted into nodes and, for each node, their *k* most similar neighbors are connected by edges. The task of clustering then is converted into detecting communities, defined as subsets of nodes where the connections among the nodes are denser within the community than connections with the rest of the graph (19). Given a computed kNN graph, Jaccard-Louvain methods employ a second step of Jaccard similarity refinement then optimizes the local structure of the kNN graph, which ultimately leads to the successful detection of communities. Jaccard similarity is defined as

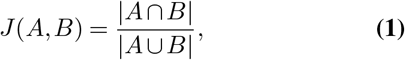

where *A* and *B* are the sets of neighbors of two cells *a* and *b*. When computing neighborhood similarities, Jaccard similarity binarizes the edge weights of the originally computed kNN graph, turning them into 1s. Therefore, the analysis only considers the presence or absence of shared neighbors, instead of focusing on the true weight of the features the cell has with its neighbors. The cells are considered more similar if they share a large portion of common neighbors, reflected by Jaccard similarity. Levine et al. (20) demonstrate that Jaccard similarity-based refinement facilitates the separation between the clusters and potentially identifies outliers from the major population on the kNN graph. Clustering algorithms making use of this strategy are collectively marked as Jaccard-Louvain methods. Scanpy, even though not using Jaccard similarity but connectivity to refine the kNN graph, is included here in a broader scope of Jaccard-Louvain methods as Scanpy is developed as an extension of PhenoGraph and Seurat.

Essentially, CosTaL adapts and extends the Jaccard-Louvain strategy. A three-step process is used by CosTaL to identify cell populations from single-cell data. First, an exact *k* nearest neighbor graph based on cosine similarity is constructed using the L2-norm k-Nearest Neighbor Graph Construction (L2knng) algorithm (21). L2knng is specifically designed to handle sparse high-dimensional data, such as single-cell profiling data, with a considerable speed advantage. The second step then involves refining the edge weights of the kNN graph using the Tanimoto coefficient, which enhances the local structure of the graph. Unlike Jaccard-Louvain methods, CosTaL uses the lengths of the original cell vectors to efficiently adjust the kNN graph weights to account for both angular and spacial separation of the points in the Euclidean space. Lastly, on the basis of the refined kNN graph, the Leiden algorithm is employed to identify communities (clusters) within the kNN graph (22). Even though CosTaL and other graph-based clustering algorithms all take similar steps and use the Leiden algorithm, how the graph is generated and refined is different. The comparisons among the algorithms are listed in Table 1.

**Table 1.** KNN-Based Clustering Algorithms

| Algorithm | Step 1. Construct kNN graph | Step 2. Refine | Step 3. Detect community |
| --- | --- | --- | --- |
| CosTaL | L2knn algorithm | Tanimoto coefficient | Leiden algorithm |
| PhenoGraph (20) | kd-tree from scikit-learn or brute force mapping | Jaccard similarity | Louvain/Leiden algorithm |
| Scanpy (23) | PyNNDescent algorithm (24) | Connectivity (25) | Leiden algorithm |
| PARC (12) | HNSW algorithm (26) | Jaccard similarity with threshold cutoffs | Leiden algorithm |

### A.1. Feature extraction

To ensure that the expression features of the cells are properly presented and compatible with the clustering algorithms, the output of the multiparametric single-cell measurements should be preprocessed prior to clustering. This process creates vector representations for the cells which are then used to construct the kNN graph.

For cytometry datasets, the features of cells denote the intensities of fluorophores (for FC) or event dual counts (for MC), which represent the amount of target proteins. The data are typically *arcsinh* transformed (*arcsinh*(*x/*5) for MC and *arcsinh*(*x/*150) for FC) to keep the readings in a linear scale (27).

For scRNA-seq datasets, the features are Unique Molecular Identifier counts, representing the absolute number of detected RNAs. In order to process scRNA-seq datasets, the most prolific preprocessing approach is the method proposed by the R package Seurat as shown in Fig. 2, including steps of ’highly variable gene selection’, ’total count normalization’, ’*log*(1 + *x*) transformation’, ’scaling’, and ’PCA’ (28, 29).

Following feature extraction, the output from cytometry and scRNA-seq differs noticeably. According to the statistics of the selected datasets presented in Table 2, cytometry datasets have fewer features than scRNA-seq datasets. In addition, the sparsity levels of cytometry datasets are generally lower than those of scRNA-seq datasets. Datasets consisting of a large number of cells (GSE110823, 1M_neurons in the table) in this category achieve sparsity of over 92%.

**Table 2.** Benchmark Dataset Statistics

| Mass/Flow cytometry datasets |  |  |  |  |  |  |
| --- | --- | --- | --- | --- | --- | --- |
| Dataset | Ref. | No. labeled cells | No. total cells | No. features | No. non-zeros | Sparsity |
| Levine_13 | (20) | 81,747 | 167,044 | 13 | 1,689,738 | 22.188% |
| Levine_32 | (20) | 104,184 | 265,627 | 32 | 6,205,110 | 26.999% |
| Samusik_01 | (9) | 53,173 | 86,864 | 39 | 1,949,437 | 42.455% |
| Samusik_all | (9) | 514,386 | 841,644 | 39 | 18,578,390 | 43.400% |
| Giordani_WT1 | (30) | 585,113 | 599,100 | 26 | 10,824,531 | 30.508% |
| Mosmann_rare | (31) | 109 | 396,460 | 15 | 5,808,969 | 2.319% |
| Nilsson_rare | (32) | 358 | 44,140 | 14 | 616,569 | 0.225% |
| scRNA-seq datasets |  |  |  |  |  |  |
| Dataset | Ref. | No. original features | No. cells | No. features | No. non-zeros | Sparsity |
| E-MTAB-3321 | (33) | 41,480 | 124 | 2,000 | 144,499 | 41.734% |
| GSE81861 | (34) | 57,241 | 561 | 2,000 | 214,688 | 80.866% |
| GSE74672 | (35) | 24,341 | 2,881 | 2,000 | 821,520 | 85.742% |
| GSE84133 | (36) | 20,125 | 3,605 | 2,000 | 792,390 | 89.010% |
| GSE110823 | (37) | 26,894 | 156,049 | 2,000 | 18,463,051 | 94.084% |
| 1M_neurons | (38) | 27,998 | 1,306,127 | 2,000 | 2,624,828,308 | 92.822% |

### A.2. Constructing the exact kNN graph using p-L2knng

Even though constructing the kNN graph to represent the similarity structure of the original datasets avoids the pitfalls of directly computing the densities in a high dimensional space, the task is still very computationally expensive, requiring *O* (*n*^2^) similarity comparisons, where *n* is the number of given cells. Many mapping algorithms resort to finding *k* nearest neighbors approximately, using one of the tree-based, hashing-based, quantization-based, or graph-based approaches (39). Nevertheless, approximate methods cannot guarantee to find the exact nearest neighbors that an exhaustive search would find, thereby undermining their reproducibility. Additionally, candidate selection and comparisons used in the approximate searching methods can be computationally expensive (40). Only a few exact search methods, such as ball-trees and KD-trees, are capable of ensuring search effectiveness while avoiding the computational difficulties associated with brute-force pairwise proximity comparisons. Still, they typically use computationally intensive pruning estimates that can not scale well with large datasets to speed up the analysis process (41, 42).

In order to tackle the scalability requirements of large-scale dataset analysis, while at the same time improving clustering effectiveness, we utilized the parallel version of the L2Knng algorithm (p-L2knng) to build an exact, rather than approximate, kNN graph (21, 43). The L2knng method efficiently uses L2-norms of feature subsets as an effective pruning strategy that avoids computation of the majority of the pairwise cosine similarities required to build the kNN graph from a sparse feature matrix. Additional details on the p-L2knng algorithm are included in Supplementary Information 1. In summary, due to multiple pruning steps inherited from L2knng and parallelization, p-L2knng significantly speeds up the kNN construction step of our framework, while at the same time constructing an exact, rather than an approximate, nearest neighbor graph.

It is worth noting that p-L2knng is designed for constructing kNN graphs using cosine similarity on non-negative input data rather than the most widely used Euclidean distance. However, as shown in earlier works, even though cosine similarity and Euclidean distance are both impacted by high dimensionality, cosine similarity typically still performs better when dealing with high-dimensional sparse data than Euclidean distance (44, 45).

### A.3. Refining the kNN graph using Tanimoto coefficient

Tanimoto coefficient, also known as Tanimoto similarity or extended Jaccard similarity/coefficient, has been broadly used in the field of cheminformatics (46), text analysis (45), and thesaurus extraction (47). Tanimoto coefficient can be calculated as

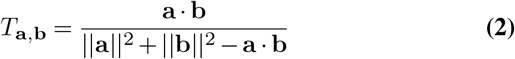

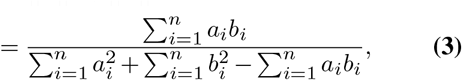

where a and b are the vector representations of two cells A and B, constructed as described in Section A.1, and *a*_*i*_ and *b*_*i*_ are the values of the *i*th component in those vector representations. Equation 3 is the typical form used for calculating the Tanimoto coefficient between two continuous n-dimensional variables. With CosTaL, the kNN graph’s edge weights are updated from cosine similarity to Tanimoto coefficient to incorporate the amplitude signal (as shown in Fig. 1).

**Fig. 1.**
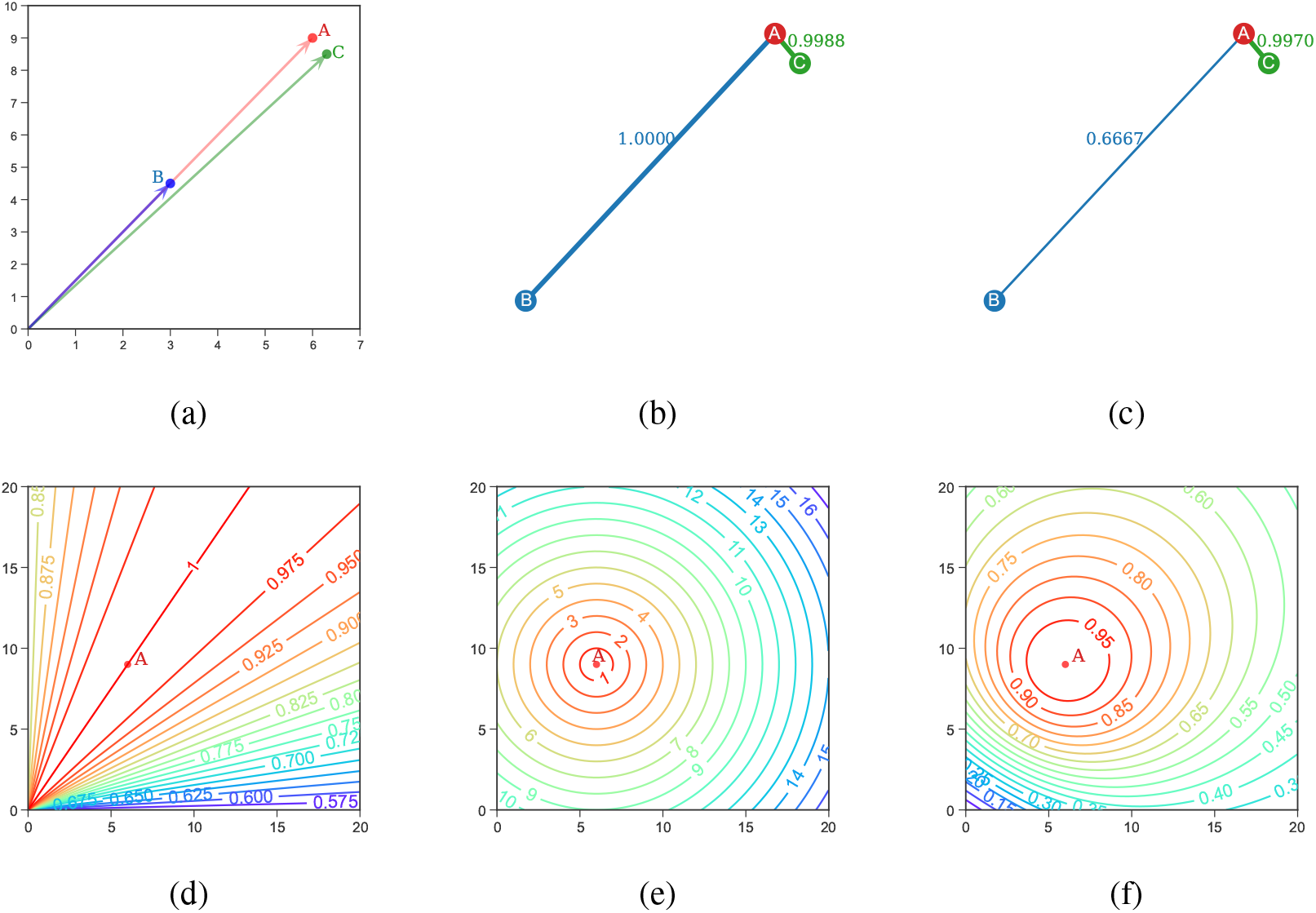
**(a), (b), (c):** Advantages of Tanimoto coefficient over cosine similarity. **(a)** Points B and C are two cells that are similar to a third cell represented by Point A. B and C are identified as A’s neighbors. **(b)** When mapping the similarity relationships with cosine similarity, the angular similarities between A&B and A&C are captured, while the dissimilarity between A&B is not reflected. **(c)** When refining the edge weights with Tanimoto coefficient, the similarity relationship is retained as no edges are removed. Moreover, it is possible to determine the amplitude difference between A and B by using Tanimoto coefficients. **(d), (e), (f):** Contour maps of a single point A’s proximity measurements in a 2D space. **(d)** Contour map of point A’s cosine similarities in a 2D space. **(e)** Contour map of point A’s neighbors mapped by Euclidean distance in a 2D space. **(f)** Contour map of point A’s Tanimoto coefficient in a 2D space. Tanimoto coefficient is able to simultaneously capture both the angular aspect of Cosine similarity and the amplitude aspect of Euclidean distance.

PhenoGraph, Seurat, and PARC all use Jaccard coefficient to refine the kNN graph. The refinement reduces spurious connections caused by outliers that share some similarities with the target but belong to a distinct population, based on the similarity of shared neighbors. Tanimoto coefficient instead focuses more on restoring the original similarity relationships among nodes, which are potentially overlooked during the initial kNN mapping using cosine similarity (Fig. 1). In Supplementary Section 2, we show how Tanimoto coefficient is an effective measure for discriminating outliers that cosine similarity would otherwise miss, being similar in this respect to the Jaccard coefficient. Aside from identifying outliers, the Tanimoto coefficient has some properties that make it a competitive similarity measure for graph refinement.

First, Jaccard coefficient refinement captures the neighborhood similarities but throws away the initial weights, which represent the proximity between the cells. On the other hand, our Tanimoto coefficient refinement includes the actual proximity measurements computed in the first step and extends them to also account for amplitude differences. As a result, CosTaL is proposing a distinctly novel strategy to construct the similarity graph needed for community detection.

Second, Tanimoto coefficient requires a smaller *k* value compared with Jaccard similarity. The Jaccard similarity algorithm treats all neighbors equally and refines the local structure of a kNN graph based on the number of shared neighbors, while the spurious links of outliers cannot be identified unless using a higher *k* value (see Supplementary Section 2 and Supplementary Fig. 1). CosTaL, on the other hand, is unaffected by this effect and requires a smaller *k* value, making it more efficient. Our experimental evaluation also confirmed this (see Section D.3).

Third, the efficiency of computing the Tanimoto coefficients is improved by reusing the cosine similarities reported by the p-L2knng algorithm, along with pre-computed L2-norms for each cell. CosTaL computes the Tanimoto coefficient as

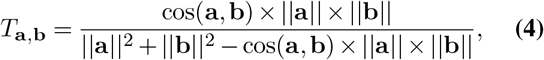

where ||a|| and ||b|| are the L2-norms of cells A and B and cos(a, b) is the cosine similarity between their vector representations.

### A.4.Community detection on the kNN graph using the Leiden algorithm

Once the network preserving the similarity information of the cell populations has been established, the cluster of cells, represented as communities on the kNN graph, can be mapped using the Leiden algorithm (22).

In the graph, a community is defined as a collection of nodes that are more densely connected with nodes in the group than with those outside the group, which can be reflected by the modularity of the graph. A detailed discussion of the computation of modularity can be found in the original article by Newman and Girvan (48). In terms of clustering, a well-clustered graph has a high modularity, which can be interpreted as an overall high degree of intra-group connections and a low degree of inter-group connections. Thus, optimizing the modularity of a graph is equivalent to clustering in such a way that the cell populations become well separated. For similarity-based kNN graphs, modularity optimization finds cell clusters with high edge weights (similarities) within clusters and low weights between clusters.

Similar to most other graph-based clustering algorithms, CosTaL employs the Leiden algorithm to optimize modularity. The Leiden algorithm is a sophisticated optimization method that uses the local move approach to merge nodes starting from singletons while optimizing graph modularity (22). As opposed to its predecessor, the Louvain algorithm, the Leiden algorithm ensures that the clusters are well-connected for the purpose of locating more high-quality clusters (49).

The advantages of modularity optimization are high effectiveness and efficiency, while a disadvantage is the resolution limit. The resolution limit of a graph with a total edge num-ber of *l* is 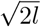, meaning that small communities with edges less than*√* 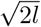 may not be found through modularity optimization (50). In light of this, some key links that represent the basic structure of the subpopulation within the kNN graph may not be well reflected in the changes in global modularity value during optimization, therefore being recognized as internal links for a greater, merged community. Since CosTaL requires smaller *k* values, the total edge number in the graph is reduced, alleviating any potential issues with the resolution limit.

Among all other algorithms, only PARC considers the resolution limit problem, and it uses aggressive truncation strategies to remove edges with weights below a threshold. Due to truncation, the network is divided into several subgraphs with a very limited number of connections. Although this is useful for detecting rare populations, it makes PARC relatively inaccurate at the global level.

### B. Benchmark datasets and preprocessing procedure

We validated the performance of CosTaL against PhenoGraph, Scanpy, and PARC in both cytometry and scRNA-seq datasets. All the data matrices were formatted to cells*×* features as the input. We picked 7 cytometry and 6 scRNA-seq benchmark datasets (Table 3) that have been used intensively in many previous comparison studies for clustering algorithms (5, 27). Based on the credibility of cell labels, we categorized datasets into three tiers. The first-tier datasets are those with labels from manual gating results (Levein_13, Levine_32, Samusik_01, and Samusik_all) or from known cell sources (E-MTAB-3321 and GSE81861). The second-tier datasets (Giordani_WT1, GSE74672, and GSE84133) are labeled with clustering algorithms with post-clustering inspections. The third group are datasets used for particular purposes. Nilsson_rare and Mosmann_rare datasets among this group were used for assessing the problem of detecting rare populations. GSE110823 and M_neurons are also in this group, primarily for evaluating the scalability of the methods.

**Table 3.** Benchmark Datasets

| Dataset | Ref. | Source | Pop. | Platform | Credibility |
| --- | --- | --- | --- | --- | --- |
| Mass/Flow cytometry datasets |  |  |  |  |  |
| Levine_13 | (20) | Human, bone marrow | 24 | CyTOF | Manual gating |
| Levine_32 | (20) | Human, bone marrow | 14 | CyTOF | Manual gating |
| Samusik_01 | (9) | Mouse, bone marrow | 24 | CyTOF | Manual gating |
| Samusik_all | (9) | Mouse, bone marrow | 24 | CyTOF | Manual gating |
| Giordani_WT1 | (30) | Mouse, skeletal muscle | 8 | CyTOF | X-Shift clustering and merge with inspections |
| Mosmann_rare | (31) | Human, peripheral blood | 1 | Flow cytometry | Manual gating |
| Nilsson_rare | (32) | Human, bone marrow | 1 | Flow cytometry | Manual gating |
| scRNA-seq datasets |  |  |  |  |  |
| E-MTAB-3321 | (33) | Mouse, Embryo | 5 | Smart-Seq2 | Sourced from 5 developmental stages |
| GSE81861 | (34) | Human, Colorectal tumor | 7 | SMARTer | Sourced from 7 cell lines |
| GSE74672 | (35) | Mice, Hypothalamus | 7 | Fluidigm C1 | BackSPIN biclustering and merge with inspections |
| GSE84133 | (36) | Human, Pancreas | 14 | inDrop | Iterative hierarchical clustering and merge with inspections |
| GSE110823 | (37) | Mouse, Brain & Spinal cord | 73 | SPLiT-seq | Jaccard-Louvain iterations and merge with inspections |
| 1M_neurons | (38) | Mouse, Brain | 60 | 10X Genomics Chromium | Jaccard-Louvain and merge with hierarchical clustering |

#### B.1. Preprocessing of cytometry datasets

Typically, cytometry datasets have some negative readings due to randomization and event calculation. Negative values are zeroed for CosTaL. As PhenoGraph, Scanpy, and PARC accept negative values, no changes were made to the data preprocessing steps of these methods. MC datasets, including Levine_13, Levine_32, Samusik_01, Samusik_all, and Giordani_WT1, were all *arcsinh*(*x/*5) transformed. FC datasets Mosmann_rare and Nilsson_rare were *arcsinh*(*x/*150) transformed (27).

#### B.2. Preprocessing of scRNA-seq datasets

For the selection of the highly variable genes, the top 2000 genes were selected for each scRNA-seq dataset as features for clustering. As required by p-L2knng, only non-negative feature values could be used as input. Since PCA transformation typically generates negative values in the Principal Components (PCs), the steps of ‘total count normalization’, ‘scaling’, and ‘PCA’ were removed in the preprocessing procedure of CosTaL as they are not needed. The illustration of the Seurat-fashioned workflow used by CosTaL versus the other algorithms is presented in Fig. 2. With CosTaL, the user not only saves more time by streamlining the preprocessing process, but also avoids having to determine how many PC dimensions should be used during the PCA transformation, which could be subjective.

**Fig. 2.**
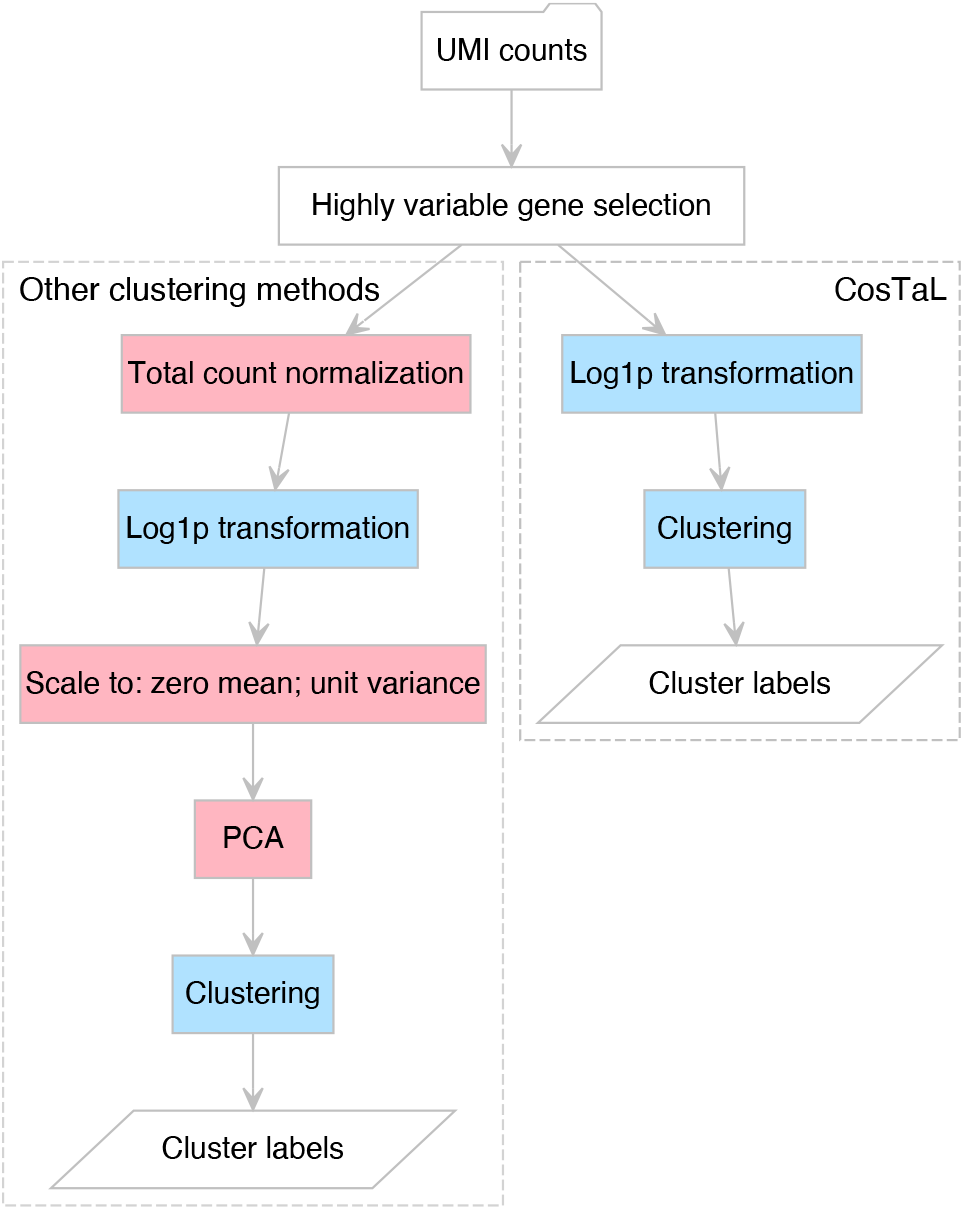
Procedures for clustering scRNA-seq data by other graph-based methods (left) and CosTaL (right). Upstream quality control (not shown) is completed to eliminate cells with insufficient information. As a result, Unique Molecular Identifiers (UMI) are generated as input, followed by selecting highly variable genes using the Seurat v3/v4 method and 2000 top genes are selected. With the unnormalized UMI matrix, a canonical preprocessing used by the other clustering algorithm involves: total count normalization, *Log*1*p* (*log*(1 + *x*), where *x* is every value in the data matrix of cells) transformation, scaling to zero mean and unit variance, and PCA transformation to reduce dimensionality. As a comparison, CosTaL only requires *Log*1*p* transformation alone. All the preprocessing steps for scRNA-seq data are executed in python using the Scanpy package.

### C. Software and Execution environment

The Scanpy package (v.1.8.2) used for preprocessing and clustering can be found at https://github.com/theislab/scanpy.git.

The PhenoGraph algorithm (v.1.5.7) can be found at https://github.com/dpeerlab/PhenoGraph.git.

The PARC algorithm (v.0.33) can be found at https://github.com/ShobiStassen/PARC.git.

The Leiden algorithm (v.0.8.2) used for community detection can be found at https://github.com/vtraag/leidenalg.git. The parameter partition method is set to *RBConf igurationV ertexP artition*, the resolution is set to 0.8, and the seed is set to its default value, which is *random*.

The p-L2Knng algorithm (v.0.2.0) can be found at http://davidanastasiu.net/software/pl2knng/. The CosTaL algorithm (v0.1.0) can be found at https://github.com/li000678/CosTaL.

Each clustering analysis on the benchmark datasets was executed on a standalone AMD ROME computing node with 64 cores and 256GB of RAM (https://www.msi.umn.edu/mangi). Clustering algorithms were executed using each of the benchmark datasets for *k* = {5, 10, 15, 20, 25, 30, 35, 40, 45, 50} and all experiments were repeated 10 times.

### D. Evaluation methods

The performance of clustering algorithms is measured by three aspects: number of clusters identified, time consumed, and effectiveness scores. While cluster number and time are straightforward, the best metric to assess clustering algorithms is yet unsettled (51). Clustering algorithms are generally evaluated using external validation methods when ground truth is available. The external validations can be divided into three different categories 1) Pair counting 2) Set overlap and 3) Information theory (52, 53). In our analysis, we took into account two popular metrics from each of the three categories in order to be comprehensive. In brief, we selected the Adjusted Rand Index (ARI) and Fowlkes-Mallows Index (FMI) measures for pair-counting-based evaluations. For overlap ratio-based evaluations, we used F-measure (F1 score). Depending on how the F1 score is harmonized as a whole for all the detected clusters, FlowCAPI F1 scores (FF1 scores) and Hungarian algorithm-based F1 scores (HF1), which have been previously described in (5, 9, 27), were adopted as our evaluation performance metrics. For information theory-based evaluations, we selected the Normalized Mutual Information (NMI) and Adjusted Mutual Information (AMI) scores. Detailed descriptions of the calculations and properties for each effectiveness score are provided in the Supplementary Information Section 3. All the scores are between 0 and 1, and 1 indicates a perfect match between the reference and the prediction (54).

## Results

### D.1. Performance comparisons on cytometry datasets using default k values

We first evaluated the performance of CosTaL (*k* = 10) against PhenoGraph (*k* = 30), Scanpy (*k* = 15), and PARC (*k* = 30). The results over cytometry datasets are shown in Fig. 3 (a) and (b).

**Fig. 3.**
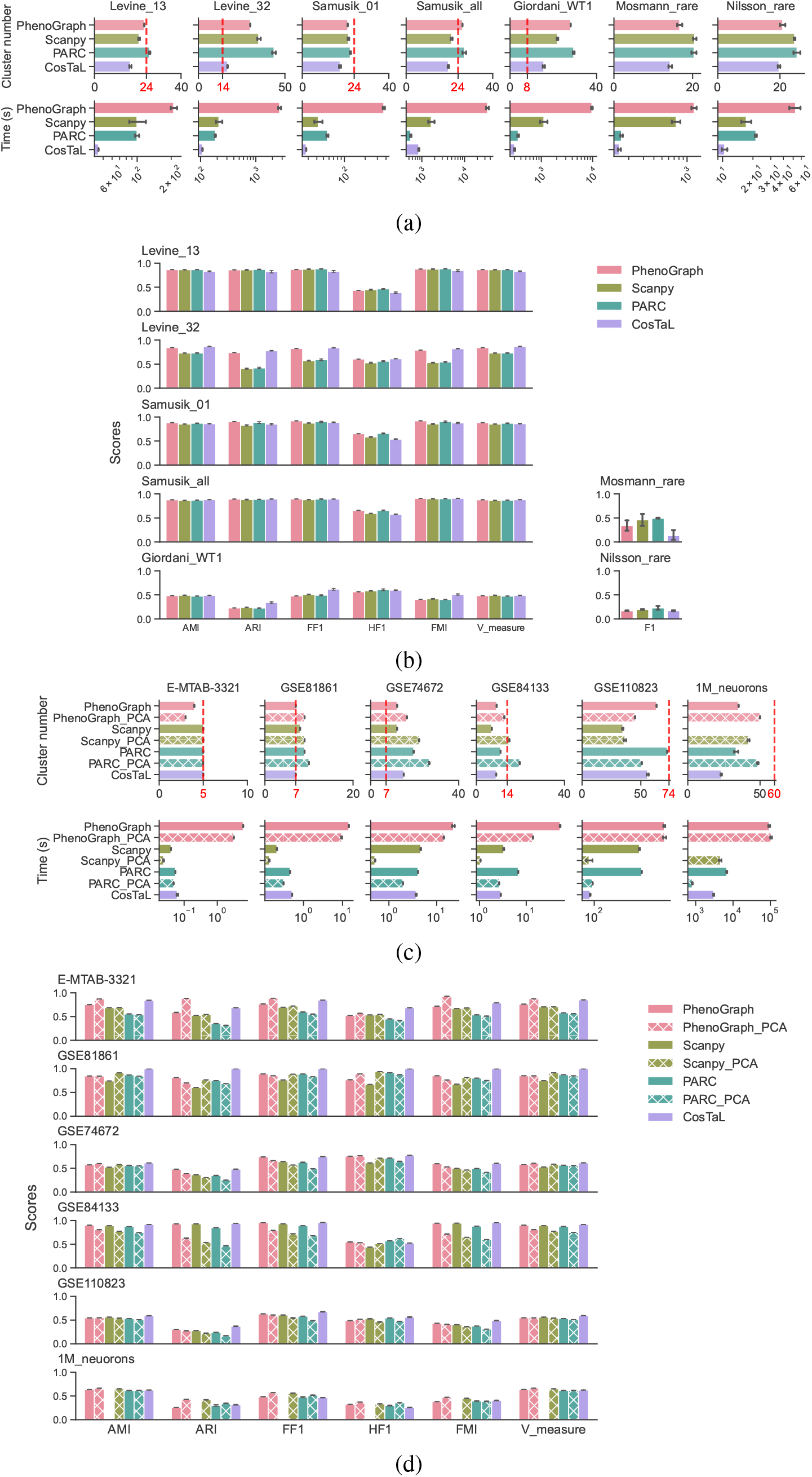
Performance of CosTaL compared with PhenoGraph, Scanpy, and PARC. **(a), (b)** Cytometry datasets. **(c), (d)** scRNA-seq datasets. **(a)** Number of identified clusters and the time consumption for each algorithm on the seven benchmark cytometry datasets. The vertical red line represents the number of cell types of the reference labels. Mosmann_rare and Nisson_rare only focus on a single rare population and the reference numbers are thus not shown. **(b)** The effectiveness scores of clustering algorithms for each dataset. AMI, ARI, FF1, HF1, FMI, and V-measure were used for all cytometry datasets except Mosmann_rare and Nisson_rare, which only used F1 score to measure effectiveness. **(c)** Number of identified clusters and the time consumption for each algorithm on the six benchmark scRNA-seq datasets. The vertical red line represents the number of cell types of the reference labels. **(d)** The effectiveness scores of clustering algorithms for each dataset. AMI, ARI, FF1, HF1, FMI, and V-measure were used for all scRNA-seq datasets. Both PCA-transformed (top 50 PCs were used) and non-PCA-transformed data were used as the input for PhenoGraph, Scanpy, and PARC. Scanpy failed in processing the non-PCA-transformed 1M_neurons dataset.

As the results show, CosTaL was the fastest among all the methods. Additionally, CosTaL generated fewer clusters than PhenoGraph, Scanpy, and PARC. The cluster number appears to affect the effectiveness scores for global-scale clustering. All six evaluation metrics show that CosTaL outperformed the other methods on the Levine_32 and Giordani_WT1 datasets, where CosTaL generated the closest cluster numbers to the references. For the Levine_13, Samusik_01, and Samusik_all datasets, CosTaL identified the least number of clusters, all below the reference number of clusters for each dataset. In these cases, CosTaL had lower HF1 scores, which require one-to-one matches between reference and clustering results, than the other methods. Yet CosTaL did not perform worse than the baselines on other performance scores.

For identifying rare populations, CosTaL was outperformed on the Mosmann_rare dataset, and none of the algorithms performed well on the Nilsson_rare dataset. The reason for this can be attributed to the fact that CosTaL identifies the fewest clusters with larger sizes, thereby affecting precision scores when calculating F1 scores. To enable the clustering algorithm to detect rare populations, CosTaL offers the option of performing iterative clustering on the identified sub-populations. In addition to the results obtained from the basic CosTaL clustering, a further level of clustering can be carried out on each cluster, which we detail in the Supplementary Information Section 4. Consequently, a more detailed partitioning of the population can be achieved, which leads to a higher F1 score. As shown in Supplementary Fig. 2, iterative two-level CosTaL clustering outperforms PARC, even though PARC is designed specifically for detecting rare populations. Overall, we can conclude that CosTaL has the advantage of efficiency while maintaining very comparable effectiveness for global-scale clustering, making it more scalable for largescale datasets. In terms of identifying rare populations, iterative CosTaL can also provide the highest level of effectiveness.

### D.2. Performance comparisons on scRNA-seq datasets using using default k value

Due to the incompatibility of CosTaL with the PCA transformation, which is generally used by other algorithms for dimensional reduction purposes in scRNA-seq datasets, we compare CosTaL with other algorithms using both PCA and non-PCA transformed data using their default *k* value. The results are shown in Fig. 3 (c) and (d).

For cluster numbers identified using CosTaL, the results are similar to those of the other methods.

In terms of efficiency, which we measure as execution time, CosTaL outperformed PhenoGraph, Scanpy, and PARC using the non-PCA-transformed data (shown as ‘PhenoGraph’, ‘Scanpy’, and ‘PARC’ in Fig. 3 (c) and (d)), but it was somewhat slower than Scanpy or PARC when they use PCA-transformed data (shown as ‘Scanpy_PCA’, and ‘PARC_PCA’ in Fig. 3 (c) and (d)). However, as we will show later, when considering both the preprocessing (including PCA) and clustering steps as a whole, CosTaL was the most efficient in most cases. Lastly, in terms of effectiveness, CosTaL performed best across all datasets with the highest AMI, ARI, FF1, FMI, and V-measure. It also had the highest HF1 scores for all but two datasets, namely GSE84133, and 1M_neurons. In this regard, CosTaL is highly competitive with other clustering algorithms.

Additionally, it is worth noting that clustering results based on PCA-transformed data were not always better than those based on non-PCA-transformed data within the baseline algorithms. As an example, PCA-transformed data always outperformed non-transformed ones on the E-MTAB-3321 dataset, while non-transformed data always outperfromed transformed data on GSE74672 and GSE84133 using all data metrics. As a means of eliminating any bias resulting from the selection of PCs, we used two midsized scRNA-seq datasets, GSE74672 and GSE84133, and evaluated the clustering performance of PhenoGraph, Scanpy, and PARC with a range of PCs from 10 to 1990. The results are shown in Supplementary Fig. 3. The results indicate that PCA did not always improve clustering effectiveness, but instead helped shorten clustering time by reducing the dimension of the features used to map the kNN graph in analyzing scRNA-seq data.

As CosTaL appears to be the most efficient method for large datasets when using the non-PCA-transformed data (Fig. 3 (c)), we further tested the overall execution time including the preprocessing steps and the clustering stages. According to the efficiency results provided in the Supplementary Fig. 4, taking into account all processing stages, CosTaL is the most efficient in this case in all but two experiments. Furthermore, Scanpy was unable to process the 1M_neuron dataset within the provided computing environment, casting doubt on its scalability as compared to the other tools. In light of these findings, CosTaL is the most appropriate choice to achieve the greatest efficiency and, at the same time, high effectiveness scores for clustering large datasets such as GSE110823 and 1M_neurons.

### D.3. Parameter influences

There are primarily two parameters that might affect the results: the similarity metrics used for mapping kNN, as well as the number of nearest neighbors (*k*) to be mapped.

The first parameter that needs attention is *k*. Considering that users have less prior knowledge, it is preferable for different *k* values to generate relatively stable results without being over- or under-partitioned. In this regard, we conducted a comparative analysis using *k* ranging from 5 to 50 for each dataset. According to the results (Fig. 4 (a)), CosTaL was capable of reaching a stable state at *k* around 10 or above but PhenoGraph and Scanpy were generally stable at *k* around 25 or above. CosTaL, therefore, required a smaller *k* value, which further reduced the amount of time required for clustering. When *k* is small, the over-partitioned clusters identified by PhenoGraph and Scanpy may not be meaningful, as evidenced by the lower effectiveness scores as well as relative studies (5, 27).

**Fig. 4.**
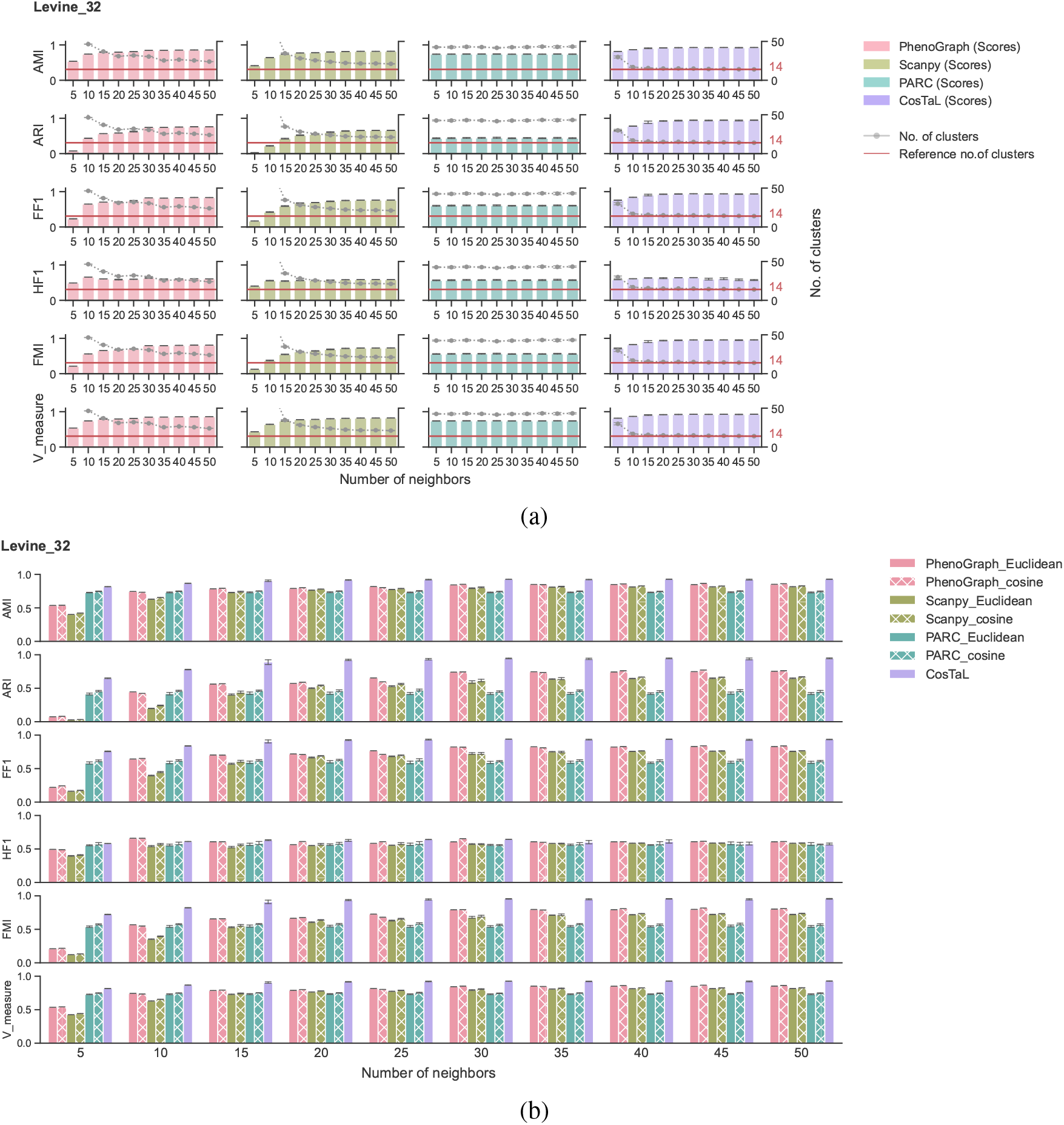
**(a)** An analysis of *k*’s effect on the clustering results of PhenoGraph, Scanpy, PARC, and CosTaL, using the Levine_32 dataset as an example. The effectiveness scores are shown as bars, and the number of identified clusters is marked by the line plot. The number of neighbors *k* ranges from 5–50. The number of cell populations in the benchmark datasets is marked with a red line as a reference. **(b)** An analysis of similarity matrices’ effects on the clustering results of PhenoGraph, Scanpy, PARC, and CosTaL, using the Levine_32 dataset as an example. Both Euclidean distance and cosine similarity were used as the proximity measures by PhenoGraph, Scanpy, and PARC, while CosTaL only used cosine similarity. The values of *k* range between 5–50.

While CosTaL only supports cosine similarity, other methods may be able to use both Euclidean distances and cosine similarity, two metrics that are most widely used for clustering. From Fig. 4(b), we can see that the choice of proximity functions can affect the results even when the same algorithm is used, though the differences are not as significant as the inter-algorithm differences. This suggests that, in determining the effectiveness of a clustering result versus a reference, the clustering algorithms are more likely to influence effectiveness than the choice of similarity metrics.

## Discussion

The current cell detection technologies are trending towards collecting information from more cells with a greater number of feature parameters. According to citation frequencies, graph-based unsupervised methods are most preferred due to their scalability, relative speed, and effectiveness (5, 10, 11, 13, 27). With the aim of improving graph-based clustering methods and making them suitable for large datasets, this report introduces CosTaL as a strategy for clustering singlecell datasets. In comparison with other algorithms, CosTaL is among the top tier in terms of speed and effectiveness with superior scalability.

Generally, CosTaL is very efficient and produces slightly underpartitioned populations based on cytometry and scRNA-seq data analysis. To measure the effectiveness, we selected six well-established methods for comparison purposes. According to the results, the CosTaL clustering algorithm has tied or outperformed other algorithms in most cases. There are basically three factors contributing to CosTaL’s efficiency. The first factor is the utilization of the p-L2knng algorithm, making the kNN mapping stage extremely fast. The second factor is the use of smaller *k* values in the kNN graph generation step, which is enabled by the Tanimoto coefficient refinement. The final factor is the shortcut of making use of the pre-computed cosine similarities provided by p-L2knng when computing Tanimoto coefficient values, which further enhances the efficiency of the refinement process.

In practice, clustering algorithms have more than a few parameters to be tuned. CosTaL simplifies the parameter tuning process, which makes it more practical for processing large datasets. The only major parameter CosTaL has to deal with is the number of nearest neighbors. We found that CosTaL could generate optimal-partitioned clusters when *k* has a value of 10 or above. As *k* increases, fewer clusters could be detected even though the change of resolution is not drastic. The method allows controlling the resolution through the parameter *k* while adding no extra effort for trial-and-error assessments since the results remain relatively stable. Specifically, for scRNA-seq data, CosTaL does not need a PCA transformation to reduce the dimensionality before clustering, further simplifying the utilization of CosTaL. Because of the efficiency of p-L2knng, CosTaL is able to map kNNs on the original feature space with 2000 highly variable genes in a short period of time, outperforming the other algorithms that require PCA transformation. Based on the clustering results, we conclude that PCA would better serve as a dimensional reduction method to ease the computational load rather than as an approach to improving the effectiveness of the clustering when dealing with scRNA-seq data. PCA is traditionally considered an effective method to reduce noise inside the sequencing data, especially for bulk analyses. However, we found that PCA does not always improve the clustering effectiveness in single-cell scenarios. This may be because most of the noise is cleaned up in the quality control and gene selection step. As a consequence, the remaining highly variable genes would be able to accurately determine the similarities among the cells without the need for PCA transformation.

Nowadays, there are many novel single-cell technologies that can monitor phenotypic and functional markers, based on either cytometry or single-cell sequencing platforms. Despite the fact that we only examined the performance of CosTaL against other algorithms in datasets of either cytometry or scRNA-seq, where the parameters are associated with proteins or transcripts, CosTaL could easily be directly extended to the analysis of other mono-modal single-cell techniques like imaging mass cytometry (55) and scATAC-seq (56).

### Key Points

CosTaL is an accurate and scalable graph-based clustering algorithm designed for analyzing single-cell data, like cytometry and scRNS-seq results. CosTaL uses p-L2knng algorithm for constructing an initial *k*-nearest neighbor graph and uses Tanimoto coefficient to refine the graph. Specifically, for scRNA-seq datasets, CosTaL does not require a PCA transformation step and still provides better scalability over largescale datasets without compromising the effectiveness of the clustering.

## Author contributions statement

Y.L. formulated the algorithm and conceived the experiments. Y.L., J.N, and D.A. developed the algorithms. Y.L., D.A., and E.A.A. wrote and reviewed the manuscript.

## ACKNOWLEDGEMENTS

This work was supported by National Institutes of Health [R01-AG020866], National Science Foundation [IIS-2002321], and the University of Minnesota [GIA University of Minnesota]. Access to research and computing facilities was provided by the Minnesota Supercomputing Institute at the University of Minnesota (https://www.msi.umn.edu).

