## Supplemental Information for "CosTaL: An Accurate and Scalable Graph-Based Clustering Algorithm for High-Dimensional Single-Cell Data Analysis"

4 CosTaL Multilevel Clustering 4

5 The Necessity of PCA Transformations 4

6 Overall Efficiency Comparison 5

### Supplementary Note 1: The L2knng and p-L2knng Algorithms

Briefly, the L2knng algorithm is divided into two phases. First, a high-quality approximate $k$-neighborhood graph is constructed very fast and then each neighborhood is improved until it has the top-$k$ nearest neighbors. Since it uses cosine similarity as the object proximity measure, L2Knng first normalizes each object vector using the L2-norm, which changes the length of the vectors without changing their angles. This preprocessing step simplifies calculations, allowing the cosine similarity of two vectors $\mathbf{a}$ and $\mathbf{b}$ to be computed as simply the dot-product of the two vectors, since

$$\cos(\mathbf{a},\mathbf{b}) = \frac{\mathbf{a}\cdot\mathbf{b}}{||\mathbf{a}|| \times ||\mathbf{b}||} = \frac{\mathbf{a}}{||\mathbf{a}||}\cdot\frac{\mathbf{b}}{||\mathbf{b}||} = \cos(\mathbf{a}',\mathbf{b}'),$$

where $\mathbf{a}' = \mathbf{a}/||\mathbf{a}||$ and $\mathbf{b}' = \mathbf{b}/||\mathbf{b}||$ are the normalized vectors.

During the first phase, the algorithm takes advantage of two principles. Since dot-products are additive and L2knng assumes vectors are non-negative, then finding vectors that have features with high values in common will lead to good neighbors. This can be achieved by using a data structure called an inverted index to investigate *candidate* vectors in reverse order of their feature value weights corresponding to the highest weight features in the *query* vector (the vector we are searching neighbors for). Additionally, once some high-quality neighbors are found, the neighborhood can be improved by looking for potential neighbors among the current neighbors' neighbors. The idea is that, given a vector $\mathbf{a}$, its current neighbor $\mathbf{b}$, and its neighbor's neighbor $\mathbf{c}$, $\mathbf{a}$ and $\mathbf{b}$ already have some high-value features in common, and $\mathbf{b}$ and $\mathbf{c}$ already have some high-value features in common. It is thus possible that $\mathbf{a}$ and $\mathbf{c}$ may have more features in common than $\mathbf{a}$ and $\mathbf{b}$, leading to a higher similarity value. These heuristic strategies lead to very quickly building an initial approximate $k$-neighborhood for each object in the dataset. In other words, each object will have $k$ neighbors but those neighbors may not be the ones with the highest similarity, i.e., the $k$-nearest neighbors.

The second phase of the algorithm seeks to find the $k$-nearest neighbors by checking as few of the remaining possible candidates as possible. This is accomplished by pruning (removing from consideration) candidates that could not theoretically meet the current minimum similarity threshold of the query object's neighborhood. In other words, given a query vector $\mathbf{q}$ whose least similar current neighbor has a similarity value of $\epsilon$, if we can show, without computing their similarity, that a candidate vector $\mathbf{c}$ cannot have a similarity value greater than $\epsilon$ with $\mathbf{q}$, then we can ignore (prune) $\mathbf{c}$. Some of the pruning is done through the use of clever data structures and specialized traversals of the vector dimensions (features). In this way, candidate vectors that have no features in common with the query (i.e., non-negative values in the same dimensions) are automatically ignored. Others are pruned by using theoretic upper bounds on dot-products, such as the Cauchy-Schwartz inequality, which states that the dot-product of two vectors is smaller or equal to the product of their L-2 norms, i.e.,

$$\mathbf{q}\cdot\mathbf{c} \leq ||\mathbf{q}|| \times ||\mathbf{c}||.$$

This is where L2knng gets its name from. At its face value, this inequality tells us nothing. Since all vectors have been normalized, we are told that all dot-products will be smaller or equal to 1. However, when we consider *feature subspaces*, this inequality becomes much more powerful. Consider, for example, that we have already computed part of the dot-product of the vectors $\mathbf{a}$ and $\mathbf{b}$ with respect to the first $p$ features in the Euclidean space, given some predefined feature

$$\begin{aligned}\cos(\mathbf{q},\mathbf{c}) &= \mathbf{q}^{\leq p}\cdot\mathbf{c}^{\leq p}+\mathbf{q}^{>p}\cdot\mathbf{c}^{>p}\\ &\leq \mathbf{q}^{\leq p}\cdot\mathbf{c}^{\leq p}+||\mathbf{q}^{>p}||\times||\mathbf{c}^{>p}||\\ &< \epsilon,\end{aligned}$$

where $\mathbf{q}^{\leq p}$ is the vector with the same values as $\mathbf{q}$ for the first $p$ features, inclusive, and all other values set to 0, and $\mathbf{q}^{>p}$ is similarly defined for the remaining features. If what we have computed so far for the dot-product plus the product of the L-2 norms of the vectors across only the remaining features is smaller than $\epsilon$, the candidate $\mathbf{c}$ cannot possibly have a higher similarity than the current neighbor with the lowest similarity $\epsilon$. Since it cannot improve the current neighborhood, it can thus be pruned. L2Knng efficiently pre-computes these partial norms and uses them to effectively prune most of the candidate vectors that would not be in the final set of $k$-nearest neighbors of the query vector. For more details on the data structures and algorithms used to quickly compute the exact $k$-nearest neighbor graph, see Anastasiu and Karypis (1). Finally, the p-L2knng algorithm (2) simplifies some of the innovations in L2Knng in order to better utilize multiple cores in a shared memory system to solve the problem in the least amount of time possible. It uses a technique called dynamic tiling to ensure the work done on each core at one time fits in the core's very fast cache memory, and a smart graph update algorithm that ensures no contention between threads when updating neighborhoods.

### Supplementary Note 2: Tanimoto Outlier Discrimination

As shown in Supplementary Fig. 1, a noise point that has connections with all $k$ neighbors is an outlier outside the major population. When refining the edge weights with Jaccard similarity, a $k$ value of 5 but not 3 can identify the outlier point. On the other hand, when using Tanimoto coefficient, a $k$ value of 3 is sufficient for discriminating the outlier point from the other points. It is evident from this toy example that both Tanimoto coefficient and Jaccard similarity are capable of discriminating outliers while Tanimoto coefficient-based refinement requires a relatively smaller $k$ value.

### Supplementary Note 3: Clustering Effectiveness Measures

**A. Pair-counting-based evaluations.** For pair-counting-based evaluations, each pair of elements (cells) is investigated to find either the agreement of both being identified in a same cluster in two partitions, or the agreement of both being assigned into different clusters in two partitions. The Rand Index (RI) corresponds to the proportion of agreement over the total number of pairs (3) and can be computed as

$$\text{RI} = \frac{TP+TN}{TP+FP+TN+FN},$$

where $TP$ is the number of pairs of elements present in the same cluster in both partitions X and Y; $FP$ is the number of pairs of elements present in the same cluster in X but not in Y; $FN$ is the number of pairs of elements present in different clusters in X but in same clusters in Y; and $TN$ is the number of pairs of elements present in different clusters in both X and Y. Adjusted Rand Index (ARI) is based on RI with a correction for chance to make sure that random assignment achieves a score close to 0 (4). It is computed as

$$\text{ARI} = \frac{RI-\mathbb{E}(RI)}{\max(RI)-\mathbb{E}(RI)},$$

where $\mathbb{E}(RI)$ is the Expected RI. The Fowlkes-Mallows Index (FMI), on the other hand, emphasizes the positive agreements, which are the pairs of elements present in the same cluster in both partitions. FMI is defined as the geometric mean of precision and recall, i.e.,

$$\text{FMI} = \sqrt{Pr\cdot Re} = \sqrt{\frac{TP}{TP+FP}\cdot\frac{TP}{TP+FN}},$$

where $Pr$ is the positive predictive rate, also known as precision, and $Re$ is the true positive rate, also known as sensitivity or recall.

**B. Overlap-based evaluations.** Set overlap-based evaluations try to match clusters with maximum overlap. Based on the overlap ratio, F-measure, also known as F1 score, is widely used and computed as the harmonic mean between precision and recall, as

$$\text{F1} = \frac{2}{\frac{1}{Pr}+\frac{1}{Re}} = \frac{2\times TP}{2\times TP+FP+FN}.$$

For assessing the effectiveness of clustering results, the F1 score is calculated for every pair of labels in the ground truth and proposed clusterings, forming a contingency table. However, an overall F-measure is desired and there are many approaches to calculate the overall F1 score. We adopted two versions that have already been used for evaluating the performance of clustering algorithms. The first version, FlowCAP-I (5), first computes the maximum F-measures of each cluster in the reference against every predicted cluster, and then aggregates them via a weighted sum normalized using the population sizes of the clusters. This enables the same predicted clusters to be matched with multiple different clusters in the reference, giving larger weight to larger populations, focusing on global clustering effectiveness. We mark F1 score calculated using this strategy as the FF1 score. The second choice is the Hungarian algorithm's version (6, 7), which calculates the overall F-measures from a strict one-to-one matching between the reference and prediction using the Hungarian algorithm to maximize the sum of F1 scores across reference populations. In this version, all clusters in the reference are treated equally. The one-to-one assignment would result in a zero F-measure in case of under-partitioned predictions, and sometimes even a merge of small clusters into larger ones would result in a drastic decrease in the overall F-measure, making it more sensitive to identify small clusters in the reference partition correctly. We

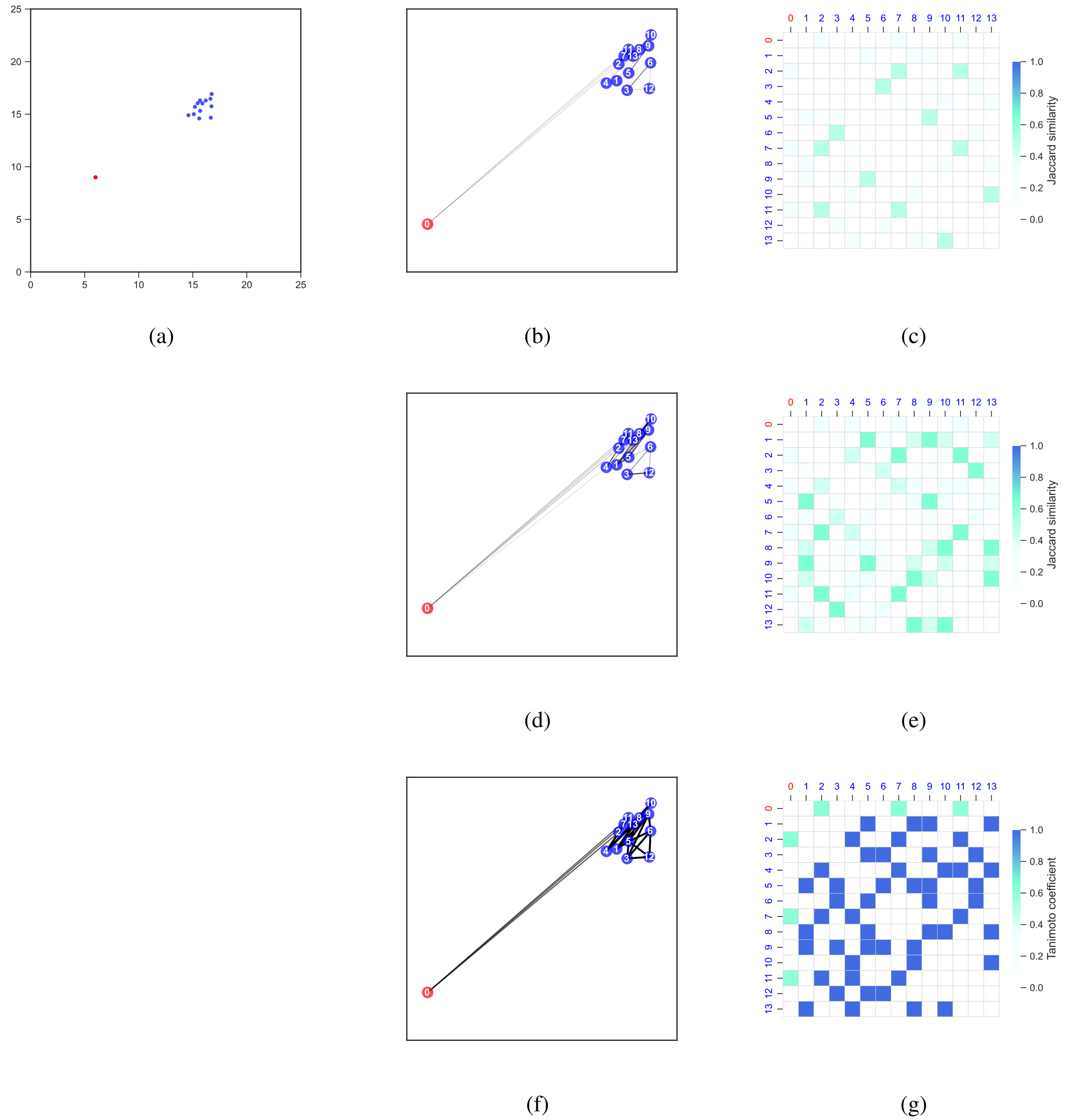

**Fig. 1. Outliers can be identified using both Tanimoto coefficients and Jaccard similarities.** **(a)** A total of 14 pseudo cells are presented in a two-dimensional feature space. Normal cells are represented by the blue colored dots, while an outlier cell is represented by the red colored dot. **(b) (d) (f)** KNN graphs for the cells in **(a)** that are constructed based on cosine similarity. **(c) (e) (g)** The edge weights of the refined kNN graphs are shown in heatmaps. The weights are averaged as kNN identifies asymmetric pairs. The heatmaps are used for assessing the effectiveness of discriminating the outlier point descriptively. **(b) (c)** Graph refined using Jaccard similarity, with $k = 3$. According to the heatmap, the outlier (No. 0) can not be distinguished from the other points. **(d) (e)** Graph refined using Jaccard similarity, with $k = 5$. According to the heatmap, the outlier (No. 0) can be distinguished from the other points. **(f) (g)** Graph refined using Tanimoto coefficient, with $k = 3$. According to the heatmap, the outlier (No. 0) can be clearly distinguished from the other points.

mark the F1 score calculated using the Hungarian algorithm as HF1. Generally, FF1 scores are less sensitive to under-partition than HF1 scores (5, 7).

The F1 score between two clusters $x_i$ and $y_j$ is computed as

$$F(x_i, y_j) = \frac{2Pr(x_i, y_j) \times Re(x_i, y_j)}{Pr(x_i, y_j) + Re(x_i, y_j)},$$

where $\mathbf{F}$ is the F1 score matrix, $Pr(x_i, y_j) = m_{ij}/|x_i|$, $Re(x_i, y_j) = m_{ij}/|y_j|$, and $m_{ij}$ is the number of matches between clusters $x_i$ and $y_j$.

Given this definition for the cluster F1 score, FF1 is computed as

$$\text{FF1} = \sum_{x_i \in X} \frac{|x_i|}{N} \max_{y_j \in Y} \{F(x_i, y_j)\}.$$

Similarly, HF1 is computed as

$$\mathrm{HF1} = \frac{\sum_{i=1}^{r}\sum_{j=1}^{s} f_{ij}a_{ij}}{\min\{r,s\}},$$

where $a_{ij}$ is from the assignment matrix $\mathbf{A}$ that has the same shape as $\mathbf{F}$, identified by the Hungarian algorithm after solving the optimization problem that minimizes

$$\underset{a}{\mathrm{argmin}} \sum_{i=1}^{r}\sum_{j=1}^{s} c_{ij}a_{ij}, \text{ s.t. } a_{ij} \in \{0,1\},$$

with $c_{ij} = 1 - f_{ij}$ and the added condition that $\sum_{i=1}^{r} a_{ij} = 1$ and $\sum_{j=1}^{s} a_{ij} = 1$, i.e., only one item is chosen per row and column of $\mathbf{A}$.

**C. Information theory-based evaluations.** Information theory can be applied to the evaluation of clustering effectiveness based on Shannon entropy, which measures uncertainty. Normalized Mutual Information (NMI) and Adjusted Mutual Information (AMI) are selected as the metrics to be used in our analysis.

In order to calculate the Mutual Information, homogeneity and completeness have to be first considered. Homogeneity is fully achieved when there is no uncertainty in inferring the true label of the clusters. Completeness is symmetrical to homogeneity, denoting the case where the predicted clusters' distributions within each true label would be completely skewed to a single predicted cluster. V-measure, being the harmonic mean between homogeneity and completeness, is a comprehensive summary of both properties to evaluate clustering performance (8, 9).

$$Homogeneity = \begin{cases} 1 & \text{if H(X,Y) = 0,} \\ 1 - \frac{H(X|Y)}{H(X)} & \text{otherwise.} \end{cases}$$

$$H(X|Y) = -\sum_{i=1}^{r}\sum_{j=1}^{s} \frac{a_{ij}}{n} \log \frac{a_{ij}}{\sum_{i=1}^{r} a_{ij}}.$$

$$H(X) = -\sum_{i=1}^{r} \frac{\sum_{j=1}^{s} a_{ij}}{n} \log \frac{\sum_{j=1}^{s} a_{ij}}{n}.$$

$$Completeness = \begin{cases} 1 & \text{if H(Y, X) = 0,} \\ 1 - \frac{H(Y|X)}{H(Y)} & \text{otherwise.} \end{cases}$$

$$H(Y|X) = -\sum_{i=1}^{r}\sum_{j=1}^{s} \frac{a_{ij}}{n} \log \frac{a_{ij}}{\sum_{j=1}^{s} a_{ij}}.$$

$$H(Y) = -\sum_{j=1}^{s} \frac{\sum_{i=1}^{r} a_{ij}}{n} \log \frac{\sum_{r=1}^{r} a_{ij}}{n}.$$

$$V-measure = 2\frac{Homogeneity \times Completeness}{Homogeneity + Completeness}.$$

Mutual Information is also based on entropy, focusing on how much uncertainty can be reduced on average to infer the cluster assignment when the label in another partition is known. Mutual information between two clusters $X$ and $Y$ is defined as

$$MI(X,Y) = \sum_{x,y} P_{XY}(x,y) \log \frac{P_{XY}(x,y)}{p_X(x)p_Y(y)}$$

where $H(X) = -\sum_{x} p_X(x) \log P_X(x)$ denotes the entropy of $X$, and $H(Y) = -\sum_{y} p_Y(y) \log P_Y(y)$ is the entropy of $Y$.

When MI is normalized by either the geometric or arithmetic mean of the two partition entropies into NMI, the effectiveness of clustering can be better interpreted since the score will be in the 0–1 range. Specifically, V-measure is equivalent to the MI score normalized by the arithmetic mean of entropies. Additionally, V-measure is equivalent to Normalized Mutual Information defined by Ana and Jain (10). The normalized mutual information is then defined as

$$\mathrm{NMI} = \frac{\mathrm{MI}(X,Y)}{mean(H(X),H(Y))},$$

where the mean function can either be the arithmetic or geometric mean. When using the arithmetic mean method, the definition matches that in Ana and Jain (10).

AMI, on the other hand, provides some adjustment to accounting for chance and excludes the expected mutual information between two random clusterings (11, 12). It is defined as

$$\mathrm{AMI}_{\mathrm{sum}}(X,Y) = \frac{MI(X,Y) - \mathbb{E}\{MI(X,Y)\}}{\frac{H(X)+H(Y)}{2} - \mathbb{E}\{MI(X,Y)\}},$$

where $\mathbb{E}\{MI(X,Y)\}$ is the expected mutual information between two random variables $X$ and $Y$.

### Supplementary Note 4: CosTaL Multilevel Clustering

In CosTaL's iterative clustering method, clustered populations are taken as input and further steps of clustering are performed to obtain more detailed partitions. Detailed segmentation facilitates the search for rare populations and results in a higher F1 score. As a precaution against over-partition, CosTaL sets a threshold (default 500) so that sub-clusters containing cells less than the threshold will not be clustered. Supplementary Fig. 2 shows the clustering results by PhenoGraph, Scanpy, PARC, CosTaL, and iterative two-level CosTaL (two levels of clustering are used) over the two flow cytometry benchmark datasets, Mosmann_rare and Nilsson_rare, in search of rare populations. As shown in the results, even though original CosTaL is not performing well, iterative CosTaL (CosTaL_iter) does yield the highest F1 scores when compared to the other graph-based clustering algorithms, at the cost of a higher running time.

### Supplementary Note 5: The Necessity of PCA Transformations

In view of the fact that CosTaL does not require Principal Component Analysis (PCA) transformation and still achieves good clustering effectiveness, we examined whether PCA transformation is necessary for clustering scRNA-seq datasets when performing clustering with the other graph-based clustering algorithms.

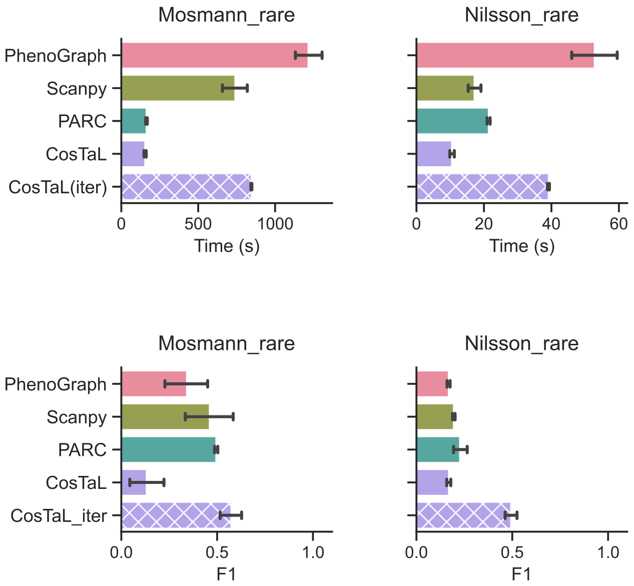

**Fig. 2. Performance of CosTaL two-level clustering option with other algorithms.**
Mosmann_rare and Nilsson_rare datasets are clustered using PhenoGraph, Scanpy, PARC, CosTaL, and CosTaL (using the iterative two-level clustering option, marked as CosTaL_iter in the figures).

We explored the relatively larger but still reliable second-tier datasets, GSE74672 and GSE84133, using different Principal Components (PCs). We tested numbers of PCs in $\{10, 20, \ldots 1990\}$. The results are shown in Supplementary Fig. 3. According to the results, all effectiveness scores of PCA-transformed input and non-PCA-transformed input were generally very similar on both datasets with all three clustering algorithms (PhenoGraph, Scanpy, PARC). In comparison to CosTaL, all three algorithms performed worse with respect to AMI, ARI, FF1, FMI, and V-measure on the GSE74672 datasets, and performed similarly to CosTaL on the GSE84133 datasets, regardless of whether PCA transformation was used or not. The results indicate that the PCA transformation does not improve the effectiveness of clustering.

### Supplementary Note 6: Overall Efficiency Comparison

We report the execution time of both preprocessing and overall clustering steps on all six scRNA-seq benchmark datasets in order to examine the scalability properties of the clustering algorithm combined with and without PCA transformation in the preprocessing step. For simplicity, the average time of 10 repeats is summarized in Supplementary Fig. 4. According to Fig. 4 (a), preprocessing with PCA requires a longer preprocessing time and is thus associated with a longer overall clustering time (as shown in Fig. 4 (b)) for PhenoGraph, Scanpy, and PARC. CosTaL generally consumes the least amount of time in the overall process. Scanpy with PCA-transformed data as the input on GSE74672 and PARC with PCA-transformed data as the input on 1M_neurons datasets are the only two cases that are faster than CosTaL. This may be attributed to the current development stage of p-L2knng algorithm as a standalone executable and its need to write files to the hard disk. A faster p-L2knng library is currently being developed that should further speed up the overall clustering time and may outperform even in these two singular cases.

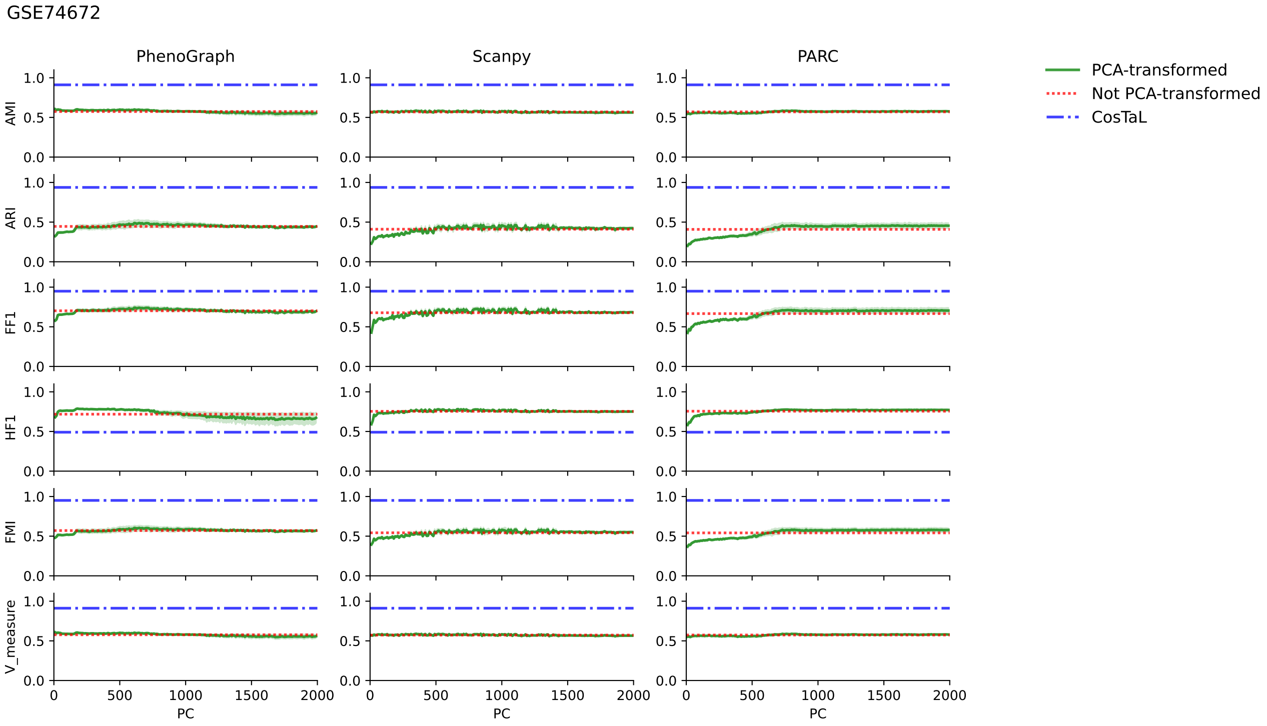

(a)

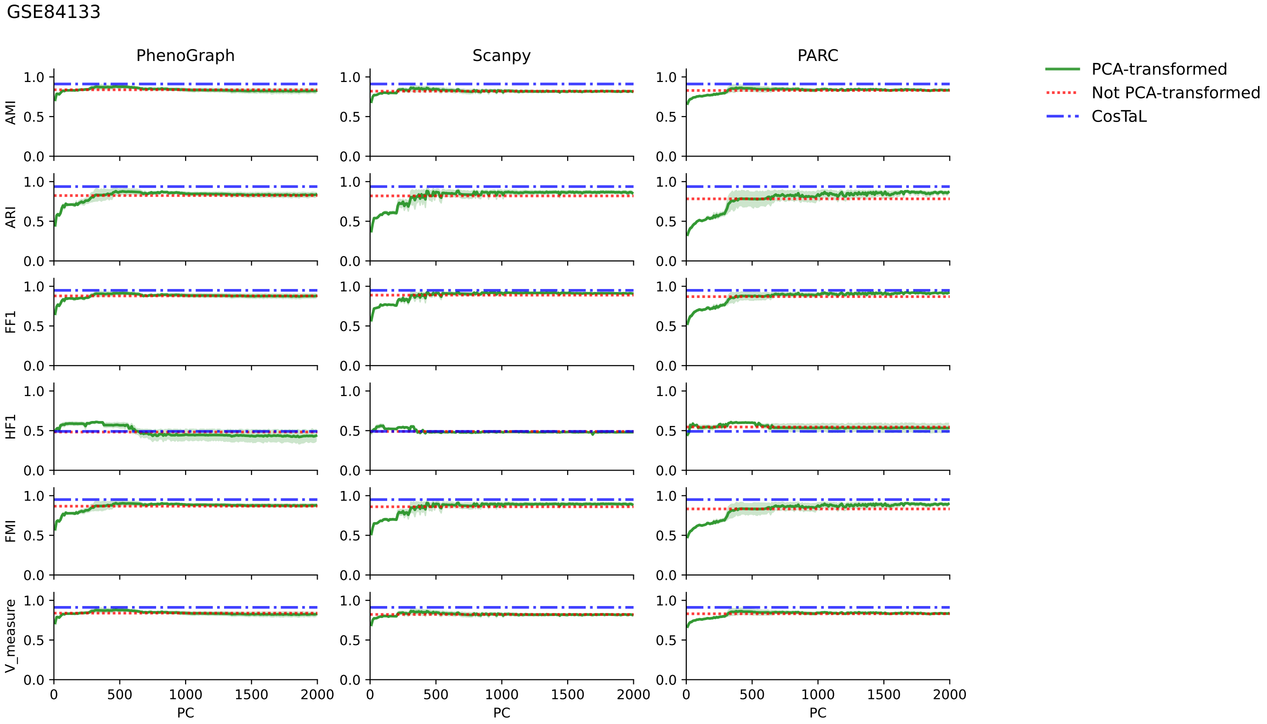

(b)

**Fig. 3. (a)** The effectiveness scores of GSE74672. **(b)** The effectiveness scores of GSE84133. **(a) (b)** Both datasets are clustered with PhenoGraph, Scanpy, and PARC. Clustering is performed using default parameters on every 10 PCs from 10 to 1990. The results of ten repeats are shown in green color in the figure, where green lines represent mean scores and light green error bands represent 99% confidence intervals. The dashed red lines indicate the mean scores of the non-PCA transformed clustering results, and the dashed blue lines indicate the mean scores of the CosTaL clustering using the default parameters repeated 10 times.

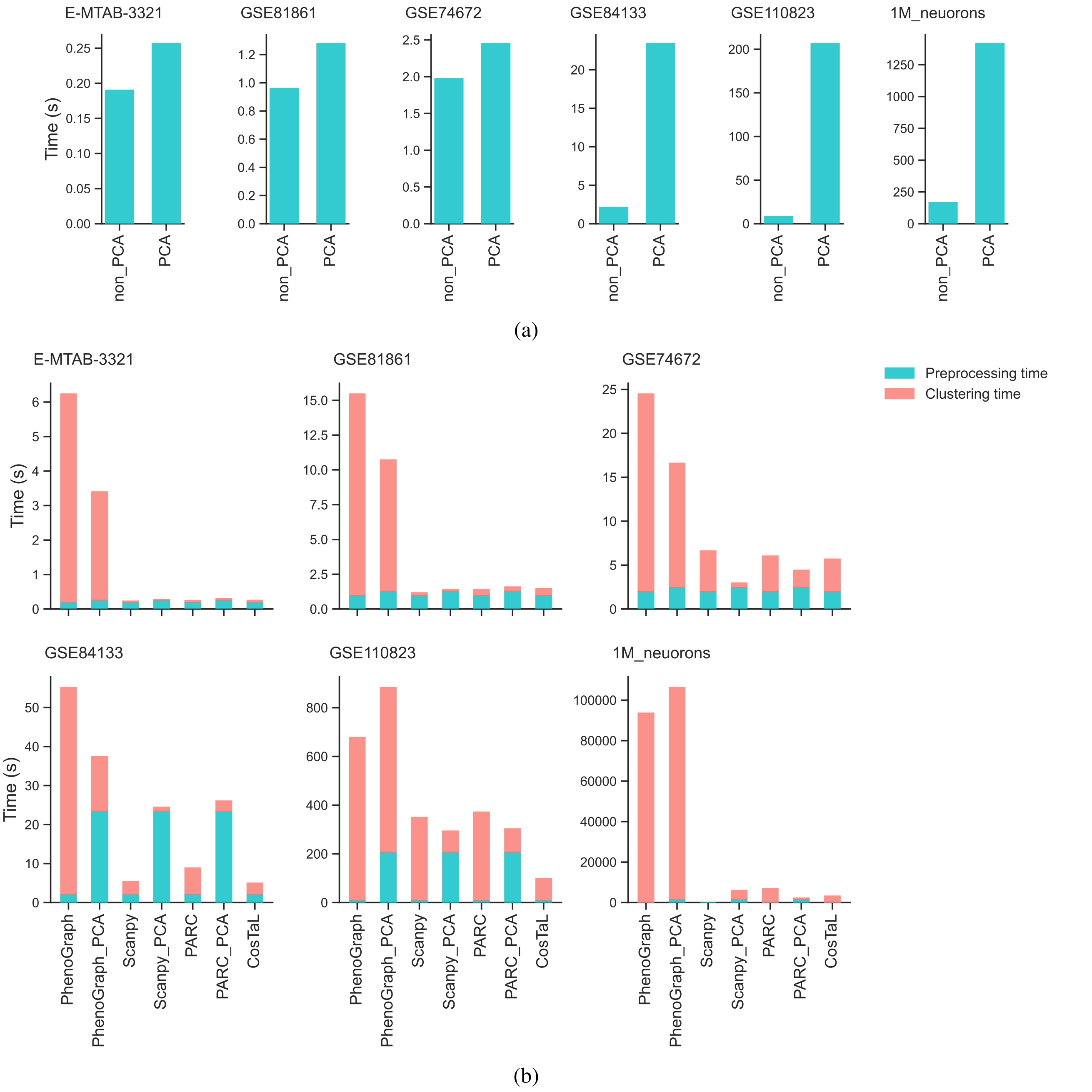

**Fig. 4. (a)** Time consumption comparisons of PCA and non-PCA transformed preprocessing steps. **(b)** Overall time consumption (including steps for both processing and clustering) of both non-PCA-transformed and PCA-transformed data as the input for clustering six scRNA-seq benchmark datasets.
